# Super-resolution STED imaging in the inner and outer whole-mount mouse retina

**DOI:** 10.1101/2022.12.18.520926

**Authors:** Leon Kremers, Kseniia Sarieva, Felix Hoffmann, Marius Ueffing, Thomas Euler, Ivana Nikić-Spiegel, Timm Schubert

## Abstract

Since its invention in 1994, super-resolution microscopy has become a popular tool for advanced imaging of biological structures, allowing visualisation of subcellular structures at a spatial scale below the diffraction limit. Thus, it is not surprising that recently, different super-resolution techniques are being applied in neuroscience, e.g. to resolve the clustering of neurotransmitter receptors and protein complex composition in presynaptic terminals. Still, the vast majority of these experiments were carried out either in cell cultures or very thin tissue sections, while there are only a few examples of super-resolution imaging in thick (> ~50 μm) biological samples. In that context, the mammalian whole-mount retina has rarely been studied with super-resolution microscopy. Here, we aimed at establishing a stimulated-emission-depletion (STED) microscopy protocol for imaging whole-mount retina. To this end, we developed sample preparation including horizontal slicing of retinal tissue, an immunolabeling protocol with STED-compatible fluorophores and optimised the STED microscope’s settings. We labelled subcellular structures in somata, dendrites, and axons of retinal ganglion cells in the inner mouse retina. Under optimal conditions, we achieved a mean lateral spatial resolution of ~120 nm (using the full width of half-maximum as a proxy) for the thinnest filamentous structures in our preparation and a resolution enhancement of two or higher compared to conventional confocal images. When combined with horizontal slicing of the retina, these settings allowed us visualisation of putative GABAergic horizontal cell synapses in the outer retina with a similar resolution. Taken together, we successfully established a STED protocol for reliable super-resolution imaging in the whole-mount mouse retina, which enables investigating, for instance, protein complex composition and cytoskeletal ultrastructure at retinal synapses in health and disease.

## Introduction

Super-resolution microscopy combines the advantages of fluorescent imaging with resolutions below the diffraction limit of light and has been abundantly used to image biological specimens [1–4]. It is especially relevant for neuroscience, where important subcellular structures (e.g. synaptic structures) have sizes below the diffraction limit [5,6]. Thus, it is not surprising that different super-resolution techniques were applied, e.g., to resolve clustering of neurotransmitter receptors [7,8] or protein complex composition in presynaptic terminals [9,10]. So far, the majority of these experiments were carried out either in cell cultures or thin tissue sections. In contrast, there are only a handful of examples of super-resolution imaging in thick specimens [11–14], as imaging deep tissue is challenging for most super-resolution techniques. The coordinate-stochastic approaches (e.g. PALM, STORM) have even more fundamental restrictions for deep imaging. In contrast, coordinate-targeted approaches, such as stimulated emission depletion (STED) microscopy, can potentially image thick specimen (e.g. the whole-mount mouse retina) by taking the advantage of optical sectioning originating from the confocal basis of the setup [15] (Figure 1A,B).

**Figure 1.**
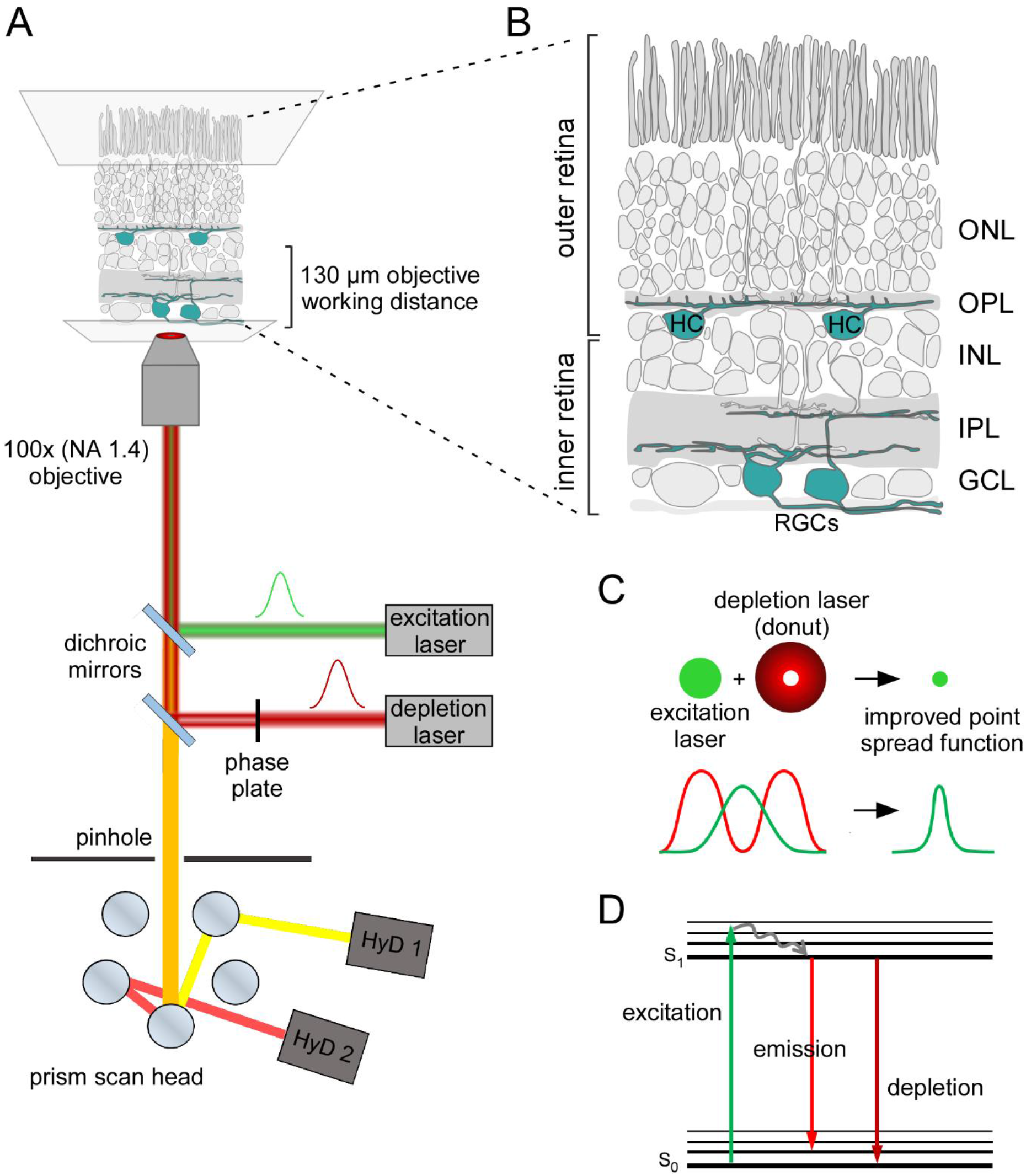
Principle of STED microscopy and application in the mouse retina. **(A)** Schematic organisation of an inverted STED microscope used in this work. In addition to the excitation laser (green) the microscope includes a red shifted depletion laser (red) which is converted into a donut shape by passing through a gradient phase plate. Depletion and excitation lasers are aligned with dichroic mirrors and focused on the sample. Emission photons (orange) are passed back through a pinhole. Distinct emission wavelengths are separated in the prism scan head and transmitted to multiple hybrid detectors (HyDs). NA, numerical aperture. (**B)** Illustration of the whole-mount mouse retina for STED imaging of retinal ganglion cells (RGCs) in the inner retina and horizontal cells (HCs) in the outer retina depicted in cyan. ONL, outer nuclear layer; OPL, outer plexiform layer; INL, inner nuclear layer; IPL, inner plexiform layer; GCL, ganglion cell layer. **(C)** Lateral resolution is improved with STED microscopy: The donut-shaped depletion laser depletes emission from the periphery of the excited volume and reduces the effective point spread function. (**D)** Jablonski diagram depicting the energetic principles behind the fluorescent depletion effect. Excitation light lifts the fluorophore from its ground state (S_0_) into a higher energetical level (excitation). The fluorophore drops into the S_1_ state via vibrational relaxation. When the fluorophore drops from its S_1_ to its S_0_ state energy is released in the form of fluorescent emission (emission). However, when the fluorophore in its S_1_ state is depleted the energy level decreases without emitting fluorescence (depletion).

STED microscopy differs from confocal microscopy by including an additional “donut-shaped” depletion laser, which is aligned to the central Gaussian-shaped excitation laser (Figure 1A,C). Here, the fluorophore is first excited by the excitation laser, and then it is either subdued by the depletion laser or spontaneously emits fluorescence (Figure 1D). The donut shape of the depletion laser enables fluorescence emission from the centre while restricting it in the spatial surround. This scales the effective point spread function (PSF) down by reducing the volume from which fluorescence is generated and detected (Figure 1C,D) [16]. The main challenge when using STED imaging lies in the additional artefacts created by thick, biological tissue: absorption, spherical aberration and light scattering intensify with increasing tissue depth [15], resulting in the generation of out-of-focus fluorescence and minimising the STED effect. To some extent, one can compensate for these effects by increasing the intensities of both excitation and depletion lasers. However, given that STED requires high laser intensities by itself, further increasing the laser power comes with a trade-off in photo-damaging effects and thermal drift as well as reducing the signal-to-noise ratio (SNR), which is already quite low in STED microscopy *per se*. Even though in-depth super-resolution imaging is challenging, there are questions in neuroscience that can be only addressed within thick specimens. For instance, STED imaging was applied in 300 μm-thick acute hippocampal slices, where it helped to resolve the fine morphology of dendritic spines upon plasticity-inducing electrophysiological stimulation [13].

As a part of the brain, the retina has a very defined layered structure [17] that enables easy access to neuronal compartments. Moreover, the chemical and electrical synapse types in the retina represent those found across the rest of the central nervous system quite well [18–20]. Given these features, valuable insights can be gained by applying super-resolution microscopy to the retina [21].

At the surface of the retinal tissue, retinal ganglion cells (RGCs) sample the visual input via the vertical photoreceptor-bipolar cell pathway and project the information via the optic nerve to higher areas of the brain (Figure 1B) [22]. Using immunolabeling against some neurofilament structures, specific types of RGCs can be easily visualised. This approach offers two advantages: first, it labels only a subfraction of all RGC types, and thus, enables the identification of individual cells. Second, labelling the neurofilament structures visualises all cellular compartments of an individual RGC – soma, dendrites, and axon. Therefore, neurofilament staining provides a suitable starting point for establishing a high-resolution imaging approach.

Horizontal cells (HCs) are interneurons in the outer retina. They have been shown to form synaptic contacts with bipolar cells (BCs) using electron microscopy [23–26]. A recent study has identified bulbs along HC dendrites, which likely represent the synaptic contacts between HCs and BCs [27]. Intriguingly, these bulbs are located below the very distal HC dendritic tips, suggesting that they do not contact cones, but co-localize with synaptic landmarks, such as mitochondria and GABA_C_ receptors. The presence of such a HC-to-BC feedforward signalling spatially separated from the HC-photoreceptor contacts is appealing, because HC feedback to photoreceptors is thought to operate locally and to be minimally influenced by global HC computations [28]. Still, HC feedforward signalling is still understudied and functional experiments testing the involvement of HC-to-BC signalling in the generation of BC responses are missing.

In this study, we developed a reliable protocol for STED imaging in the whole-mount mouse retina. First, we optimised the sample preparation and imaging settings for imaging RGC neurofilament structures close to the surface of the retinal tissue. Second, we adapted our approach for imaging synaptic structures of HCs deeper in the outer retina.

## Methods

### Animals & whole-mount retina tissue preparation

Retinae from adult (4-13 weeks old) male and female mice of the C57BL/6J wildtype line were used for this study. The animals were deeply anaesthetized with isoflurane (CP-Pharma, Germany) and were sacrificed by cervical dislocation. All animals were handled in accordance with the European and national government regulations following the European animal welfare law. The eyes were quickly enucleated, and all further dissection steps were performed in 0.1 M phosphate saline buffer (PBS) (pH 7.4). Cornea, lens and vitreous body were carefully removed. The retina was dissociated from the eyecup and mounted RGC side-up as a whole retina or cut in three or four pieces on black nitrocellulose membrane (0.8 mm pore size, Millipore, Ireland).

### Immunohistochemistry of the inner retina

The whole-mount retina preparations (for imaging of the inner retina) were fixed using 4% paraformaldehyde (PFA) in 0.1 M PBS for 20 minutes at 4°C, washed with 0.1 M PBS (6 x 20 minutes at 4°C) and blocked with blocking solution (10% normal goat serum (NGS) and 0.3% Triton X-100 in 0.1 M PBS) overnight at 4°C. Afterwards, the samples were incubated with primary antibodies (see Table 1 below) solution with 0.3% Triton X-100 and 5% NGS in 0.1 M PBS for 4-9 days at 4°C. The samples were then washed with 0.1 M PBS (6 x 20 minutes at 4°C) and incubated with secondary antibody (see Table 2 below) solution in 0.1 M PBS overnight at 4°C. After another washing step (6 x 20 minutes at 4°C), the retinae were embedded in mounting media and mounted between the slide and the high precision coverslip (Marienfeld, Precision cover glasses, No. 1.5H, Germany). Different types of mounting media were used. ProLong Gold (ThermoFisher Scientific, USA) and Vectashield (Vector Laboratories, USA) were used according to the Manufacturers’ protocol. Abberior TDE Mounting Medium O (Abberior, Germany,) was used according to manufacturer prescriptions with elongation of each incubation step to 1 hour (see description below). The samples were covered with high-precision coverslips (No. 1.5H, Carl-Roth GmbH, Germany) and sealed with transparent nail polish and left overnight at 4°C.

**Table 1.**
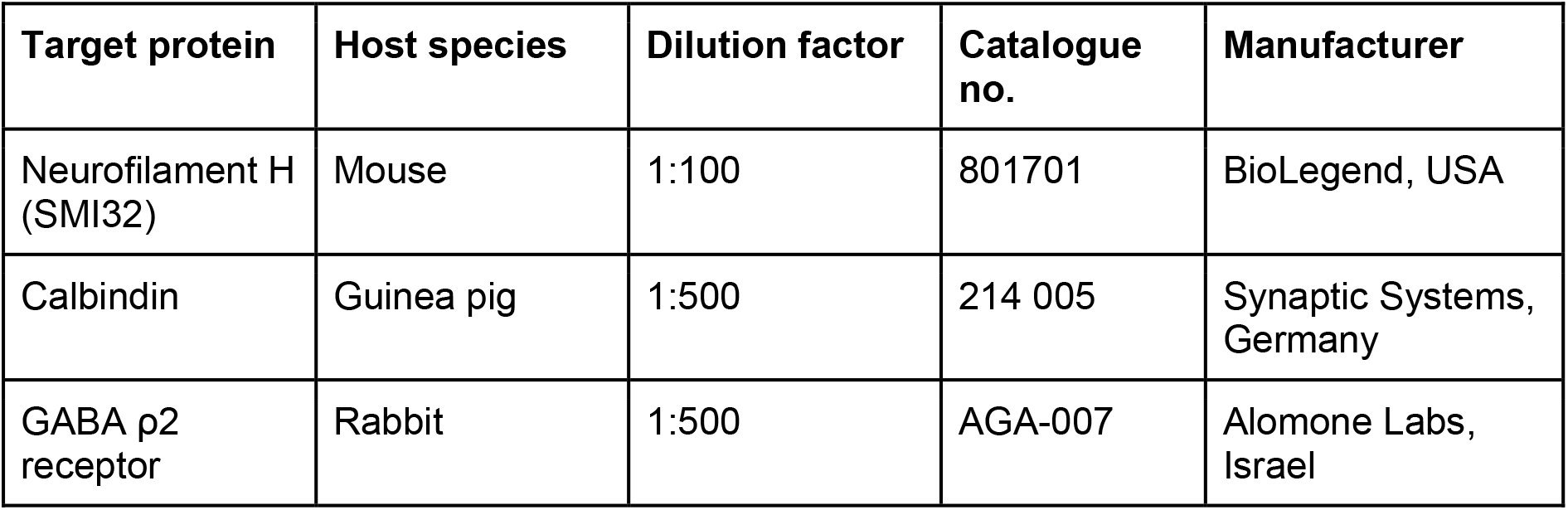
The following primary antibodies were used:

**Table 2.**
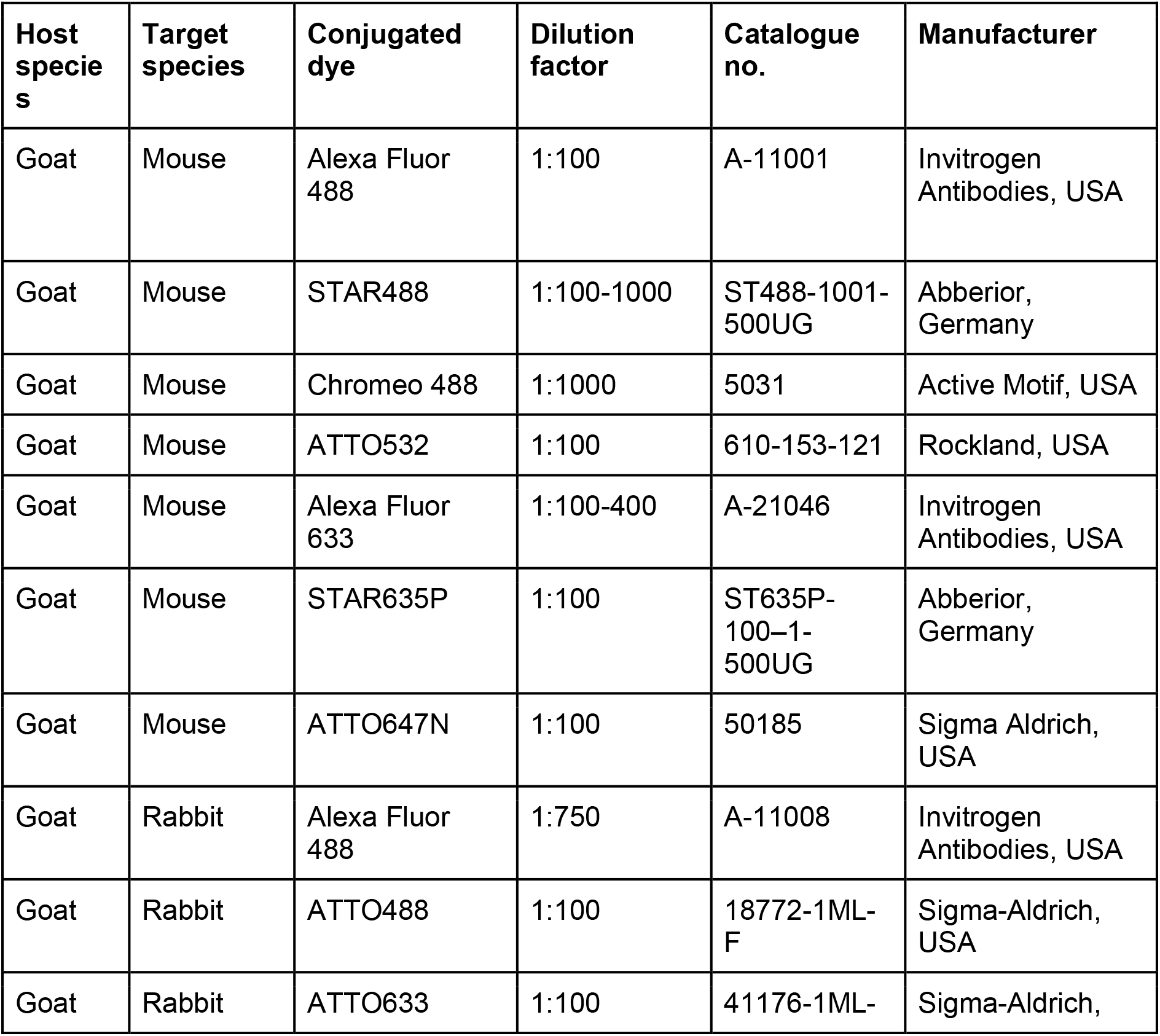
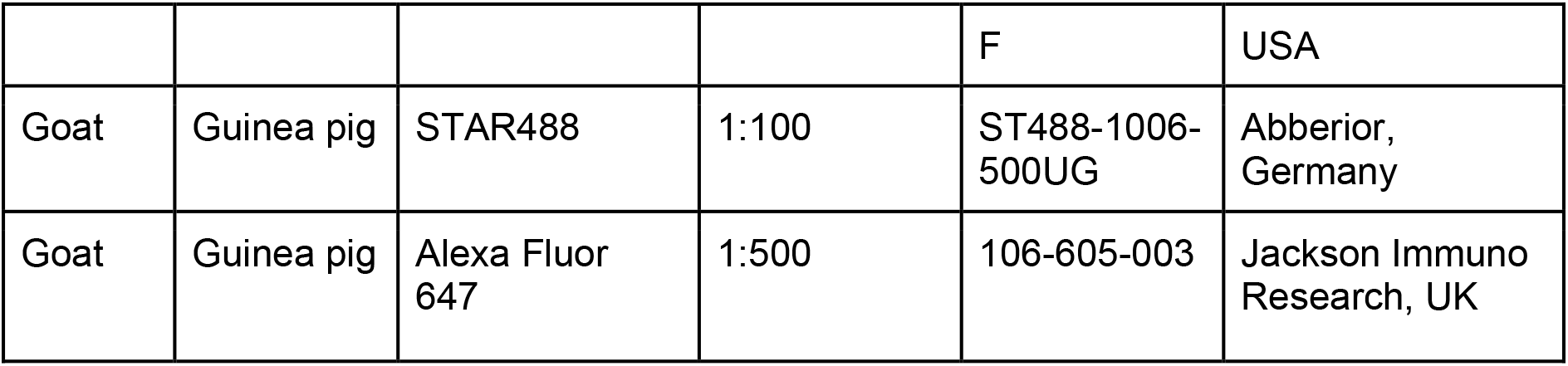
The following species-specific secondary antibodies were used:

### Cryosectioning and immunolabeling of horizontal sections of the outer retina

To prepare retinal pieces for horizontal sectioning (and imaging of the outer retina), retinal pieces, isolated as previously described, were fixated in 4% PFA solution for 20 min as described above. After fixation, the retinal pieces were washed in 0.1 M PBS (3 x 10 min) at 4°C, separated from the cellulose membrane and passed through a series of incubations in sucrose-PBS-solutions with increasing concentration. All sucrose incubations were performed at 4°C. First, retinae were kept in 10% sucrose solution for 1 to 2 h, until all pieces sank to the bottom. Retinae were then transferred into 20% sucrose solution and incubated for an additional 1 to 2 h, again until all pieces subsided, before being transferred to a 30% sucrose solution, in which they were kept overnight. Retinal pieces were transferred into tissue freezing medium for preincubation before being mounted to the sample holders of an Epredia CryoStar NX50 Cryotome (Thermo Fisher Scientific, USA). In order to section the retinae along the xy axis as horizontally as possible, tissue freezing medium was applied to the sample holders of the cryotome and frozen solid. The cryotome was then used to cut an even plane into the frozen medium big enough to fit one retinal piece. Retinal pieces were aligned on a glass slide wrapped in parafilm with the RGC layer facing downwards. Single pieces were then picked up by carefully descending the plane of frozen medium with the sample holder on it. Additional freezing medium was used to fully cover the retina, before the sample was quickly frozen with liquid nitrogen. The sample holder was then placed back into the cryotome and 50 μm thick horizontal sections of the retina were cut. Cut sections were picked up using superfrost slides (Thermo Fisher Scientific, USA) and dried for 1 h at 37°C on a heating plate.

Retinal slices mounted on microscopy slides were surrounded by 120 μm thick silicon spacers (Secure-seal spacer, Invitrogen, USA) or PAP pen (Science Services, Germany) and solutions were directly applied on the slides. After cryosectioning, the retinal sections were washed in 0.1 M PBS (6 x 20 min) at 4°C. 0.1 M PBS was removed and unspecific antigens were blocked by incubating the retinae in 10% NGS and 0.3% Triton-X-100 in 0.1 M PBS overnight. The blocking solution was then removed, and primary antibodies were added (see Table 1). The antibodies were diluted in 5% NGS and 0.3% Triton-X-100 in 0.1 M PBS to the concentration specified by the respective manufacturer. To accommodate the distinct diffusion time of the antibody within the cut samples of minor thickness, retinal slices were incubated for two days. After incubation, unbound antibodies were removed by washing the retinae again (6 x 20 min) with 0.1 M PBS at 4°C before adding the secondary antibody. Secondary antibodies, which are conjugated to fluorophores of choice (see Table 2), were diluted in 0.1 M PBS to the concentration specified by the manufacturer and retinae were incubated with the solution overnight. To remove excess antibodies, the retinae were again washed in 0.1 M PBS (6 x 20 min) at 4°C. All retinae were mounted with a 2,2’-thiodiethanol-based embedding medium (TDE, Abberior, Germany. Some retinae were mounted with the 120 μm thick spacers between slice and coverslip. One drop of 10% TDE solution was added to cover the retinal pieces and left to incubate for 1 h at 4°C. Afterwards, the 10% solution was exchanged with 25% solution and incubated again for 1 h, before being exchanged again for a 39% TDE solution. The retinae were incubated in the 39% solution for 45 min before the medium was substituted for the final 97% TDE solution. The samples were left to incubate for an additional 45 min before and high-precision coverslips (No. 1.5H, Carl-Roth GmbH, Germany) were carefully put on top of the retinal pieces and spacers. The sample was sealed by applying nail polish to the edges of the coverslip and the polish was left to dry overnight at 4°C before imaging experiments were performed.

### Confocal and STED imaging

Confocal and STED imaging were both performed at the same microscope setup using a DMi8 inverse microscope (Leica, Germany) with three oil immersion objectives with 20x, 63x and 100x magnification (Leica, Germany) in combination with the TCS SP8 STED setup (Leica, Germany). The microscopy setup included three pulsed excitation lasers, one with 488 nm (Leica, Germany) and two additional lasers with 532 nm and 635 nm wavelengths (OneFive, Switzerland). For depletion, a continuous-wave 592 nm and a gated/pulsed 775 nm STED laser were used. The intensities of the depletion lasers were experimentally chosen and were 0.65 W (43% of maximum value) for the 592 nm laser and 0.45 W (30% of maximum value) for the 775 nm laser. The microscope function was controlled via a connected computer using LasX software (Leica, Germany). Fluorescent emission was split and quantified using the integrated prism scan head and photomultiplier tubes (PMTs) and/or hybrid detectors (HyDs). For STED imaging only HyDs were used. The scan head allowed for the free selection of the emission wavelength spectra to be captured and measured. For STED imaging, xy-pixel and z-step size were chosen in accordance with the Nyquist–Shannon sampling theorem. This meant that STED imaging was only performed using the 100x objective and using an image size of 2048 x 2048 pixels. An additional 3-4 x zoom was applied, resulting in an effective pixel size of 14 - 20 x 14 - 20 nm. Minimal z-step size was manually calculated and set to 130 nm. Bidirectional scanning was enabled, and each line was scanned three times with pixel intensities being accumulated. For each z-section of an image stack three lines or frames were imaged and intensity values averaged. Excitation and depletion laser intensities as well as PMT/HyD gain were determined via testing of signal strength and photobleaching. Laser intensities and gain thus differed on a case-to-case basis but were kept consistent within experiments. Confocal imaging was performed by disabling the depletion lasers while keeping the excitation lasers on. If the same region was imaged with both STED and confocal microscopy, confocal imaging was typically performed before STED to prevent photobleaching from the high intensity depletion laser. When the same region was imaged in multiple fluorescent channels, fluorophores with higher excitation/emission wavelengths were imaged first. Imaging data was saved as lif-files, with z-stacks being represented as different series within one file.

### Image processing and analysis

Acquired images were processed and analysed by LasX (Leica Microsystems) and ImageJ (National Institutes of Health, USA) software. Stabilisation and deconvolution of STED images were performed by Huygens Software (Version 17.10.0p6 64b, SVI, Netherlands). The deconvolution was performed with the Classical Maximum Likelihood Estimation (CMLE) algorithm under experimentally defined settings. Images were deconvoluted using theoretical PSFs calculated for the DMi8 STED microscope and the 100x (NA 1.4) objective used in this work. No lateral drift and bleaching corrections were performed, and most settings were kept at automatically defined values. The background level was estimated by software, the quality threshold was 0.001, the number of iterations was 50, the SNR was set to 7 for STED images. In ImageJ, the original and deconvolved z-stacks were typically transformed into a single maximum intensity projection and saved as raw files for further analysis. For presentational purposes image contrast and brightness were automatically optimised, channels depicted in defined colours and scale bars inserted. All quantifications and processing were performed in unprocessed (= not deconvolved data) unless otherwise specified (see Figure 8).

Image analysis was performed using the open-source software ImageJ in its Fiji distribution and custom scripts written in the R programming language. For direct extraction of intensity values, a line selection was performed in ImageJ and values along the selection were extracted for all channels manually or using a custom written ImageJ macro. For determination of the full width at half maximum (FWHM), line profiles were fitted (Gaussian curve, non-linear least square (NLS) approximation) using R free programming software. In short, raw image files were imported into R and transformed into intensity value matrices. Coordinates were selected using the shiny plug-in for R and intensity values along a vector between both coordinates were saved. A Gaussian curve was fitted to these values using the NLS approach and the nls-multstart package. The general formula for the fitted curve was defined as:

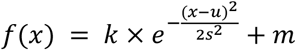

The x values with *f*(*x*) = *m* + 0.5 × (*f_max_* – *m*) were defined and the distance between both x values was calculated as the FWHM: For structures in the inner retina (RGC structures), we used an unconstrained NLS fit. In the outer retina (HC structures and GABA receptors), we used multiple constraints in our model: First, we defined *s* as *s* ≥ 25 determining the width of the Gaussian curve with a minimal FWHM of 58.8 nm). With our imaging system and sample probes we did not expect a resolution below this limit. Second, under the assumption that the brightest pixels along the line represent the structure we are interested in, we defined *f_max_* ≥ maximum intensity, and thus, prevented the model from being biassed by unspecific background noise. Third, we defined *u* as 50 ≤ *u* ≤ length of line - 50 nm to ensure that the model doesn’t fit the curve to unspecific noise at the borders of the extracted vector. Overall, constraining our model produced fits that improved in describing the observed signal.The process was repeated with the previously defined coordinates for all corresponding images. Ratios between corresponding confocal and STED FWHMs as well as STED and deconvolved STED FWHMs were calculated and saved along the absolute FWHMs, the formulae of the fitted curves, the extracted intensity values and the determined coordinates in an Excel (.xlsx) file. For the resolution enhancement factor (REF), the normalised pixel intensities of Calbindin and SMI32 stainings were subtracted for the line defined across the bulb/non-bulb.

### Statistics

Data representation and statistical analysis was performed using Prism 9 software (GraphPad, US) or in the R programming language. For Prism 9, data sets were copied into grouped tables as required and statistics were calculated using the analysis function. For complex data featuring various groups, each with multiple subcategories, 2-way ANOVA combined with a Tukey’s multiple comparisons test was performed. Paired data with only two categories was subjected to Shapiro-Wilk normality testing and two-tailed paired t-test was used for analysis of normally distributed data, Wilcoxon signed-rank test was used otherwise. Unpaired data with only two categories was analysed using an unpaired t-test or Wilcoxon rank sum test. Significance was defined as: p > 0.05 = ns, p < 0.05 = *, p < 0.01 = **, p < 0.001 = ***, p < 0.0001 = ****, p < 0.00001 = *****, p < 0.000001 = ******. Mean values in text and figures are given as mean ± standard error of the mean (SEM).

## Results

To establish STED microscopy (Figure 1A,B) in the mouse retina, we used whole-mount preparations labelled with a full-sized primary antibody against neurofilament H (from here on: SMI32 labelling) and secondary antibodies conjugated with synthetic dyes. The required labelling density was achieved by increasing the concentrations of both primary (2x, Table 1) and secondary (≤ 10x, Table 2) antibodies compared to commonly used concentrations of respective antibodies. We improved the sample preparation by choosing the best-performing spacer type between slide and coverslip, mounting medium, and synthetic dyes (see Methods). We tried both conventional dyes (Alexa Fluor) and new-generation dyes (STAR, ATTO, Chromeo) specifically developed for STED microscopy. We tested photostability and selected dyes with the lowest bleaching effects at 488 and 635 nm excitation. For both green and far-red dyes, new-generation dyes were more photostable than Alexa Fluor dyes. For far-red dyes (ATTO647N and STAR635P), we never observed bleaching with our experimental conditions.

To avoid physical squeezing of the retinal tissue upon mounting, we placed spacers in between the slide and coverslip. For this purpose, we used commercially available silicon spacers.

The choice of mounting medium was determined by its refractive index (n). We tested four different mounting media: Abberior Liquid AntiFade (n = 1.38), Vectashield (n = 1.47), Abberior TDE (n = 1.51) and ProLong Gold (n = 1.47). Abberior Liquid AntiFade had a lower refractive index than the immersion oil (n = 1.52) and was thus excluded from further experiments. We hypothesised that ProLong Gold should be the most stable medium as it was the only polymerizing mounting medium in our study. However, we observed strong axial drift when switching from confocal to STED mode, which we could not correct for. Vectashield mounting medium is not recommended by STED manufacturers because it absorbs light in the red range of the spectrum and is incompatible with large-Stokes shift dyes. However, in our experiments, only a minor difference in resolution with Vectashield and Abberior TDE could be observed (resolution of STED images determined as the full width at half maximum (FWHM, see Methods); with Abberior TDE: 118.2 ± 4.7 nm (n = 24 structures, n = 3 images); with Vectashield: 144.9 ± 7.6 nm (n = 20 structures, n = 3 images; p = 0.01, Wilcoxon rank sum test). Finally, we chose Abberior TDE as the most stably performing mounting medium with its only constraint being the short lifetime of the specimens at around 1.5 - 2 weeks in our hands.

### Optimization of microscope settings for STED microscopy

The main feature of a scanning STED microscope is the depletion laser (Figure 1C,D). Our STED microscope was equipped with two depletion laser lines with 592 and 775 nm wavelengths. Whereas the 592 nm depletion laser was used together with the 488 nm excitation laser, the 775 nm depletion laser depleted the emission of fluorophores excited with either the 532 or 633 nm excitation laser. The orange beam (592 nm) is a continuous-wave laser, whereas the far-red one (775 nm) is a pulsed/gated laser (Figure 2B). The resolution of STED imaging depends – among other factors – on the saturation factor (I_max_/I_s_, Figure 2A). With the 592 nm continuous-wave depletion laser, we could achieve a maximum saturation factor of 7.5 without severe bleaching of the 488 nm dye at laser intensity 0.65 W (Figure 2C). In contrast, with the 775 nm laser the saturation factor was as high as 30 at relatively low laser intensity of 0.45 W (Figure 2D), likely due to the fact, that a high density of 775 nm photons is ‘pumped’ into the pulsed events whereas the photon number in the between-pulse intervals is minimal (Figure 2B). Therefore, we suggest that using the pulsed depletion laser results in both better resolution and minor photo-damaging compared with the continuous wave depletion laser.

**Figure 2.**
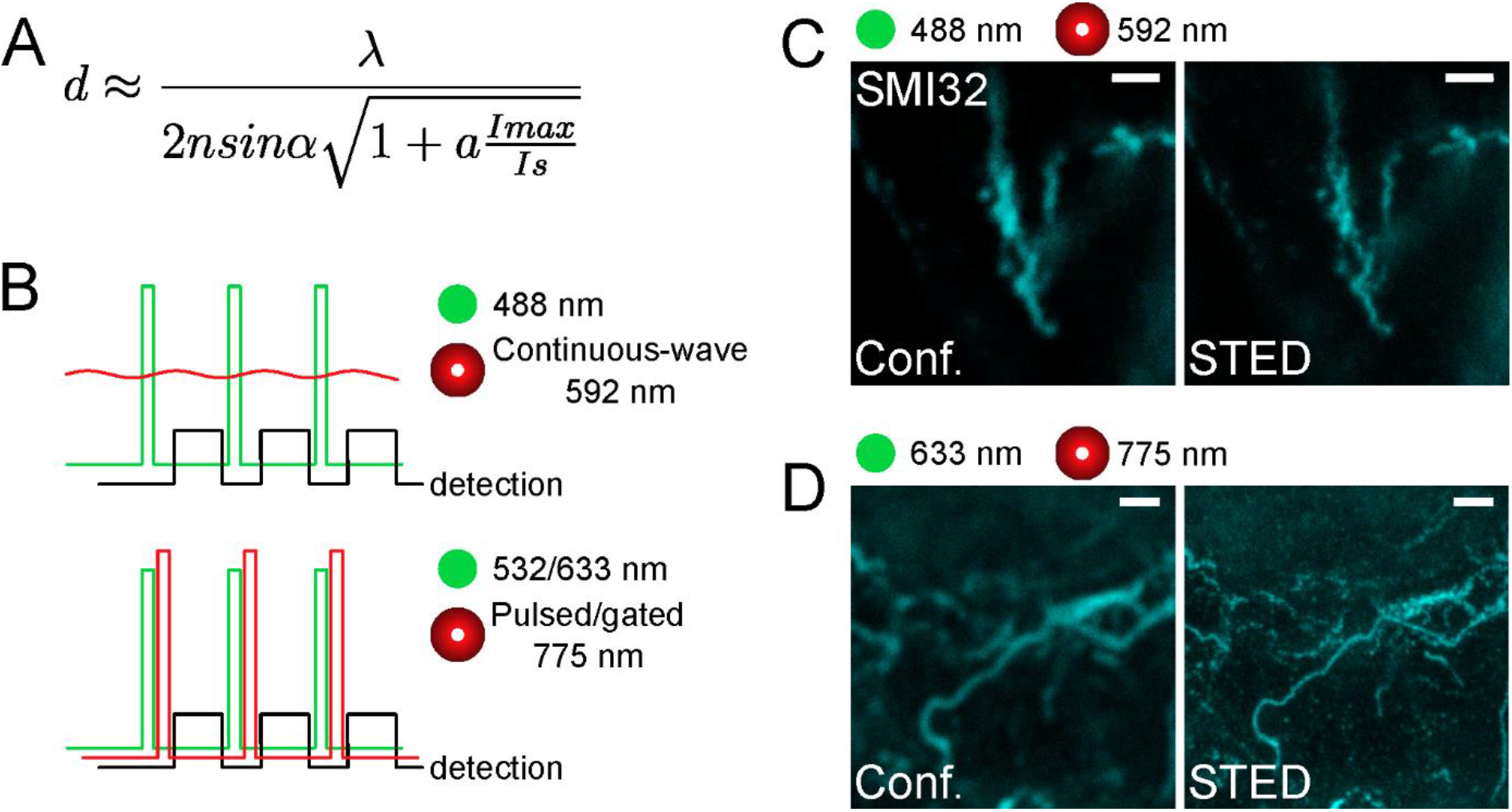
Representative SMI32-labelled structures in the inner retina acquired with different STED lasers. **(A)** Formula for the lateral resolution of STED microscope with the saturation factor I_max_/I_s_ with Imax as the maximally applied laser intensity and I_s_ as the STED laser intensity at which half of the fluorescence is lost. (**B)** Temporal conditions of STED imaging. Ideally, all depleting photons act when fluorophores are in the singlet-excited state S_1_ and fluorescence must be registered after the stimulating photon’s action. Experimental time sequences are shown for the excitation (green), the depletion (red), and the emission signal detection (black) for continuous-wave STED (592 nm, top) and gated/pulsed STED (775 nm, bottom) microscopy. (**C,D)** Representative confocal (left) and STED (right) images acquired with different excitation and STED depletion lasers (592 nm in C, and 775 nm in D). For C, the excitation wavelength was 488 nm, and the dye was STAR488. For D, the excitation wavelength was 635 nm, and the dye was ATTO647N. Scale bars: 1 μm.

In general, we obtained more comparable fluorescence intensities by adjusting the excitation laser intensity for every specimen separately and increasing it with larger imaging depths. We defined the xy-pixel size as 14 - 20 nm according to the Nyquist-Shannon sampling theorem. Other imaging settings were adjusted experimentally. The bit depth was 12 bit, and we used 3x line accumulation with or without 3x frame averaging. Frame averaging was commonly used to decrease unspecific noise, however, in our case, the laser exposure was sometimes too high and led to thermal effects and physical deterioration of the sample upon imaging from the same focal plane.

### Super-resolution microscopy of retinal ganglion cell structures in the inner retina

Super-resolution STED microscopy of thick specimens remains challenging due to scattering within biological tissue, which leads to decreased depletion efficiency in deeper layers. This restricts efficient STED imaging to the superficial 50-70 μm of the sample. In the retina this corresponds to the ganglion cell layer with the somata and axons of RGCs, and the inner plexiform layer, which roughly comprises of dendrites and synaptic connections of RGCs, BCs and amacrine cells (ACs) (Figures 1B,3A). SMI32 labelling reveals intermediate filaments in axonal bundles of RGCs (Figure 3B) as well as dense cytoskeletal network structures in RGC somata (Figure 3C) at a depth of 25-30 μm. Imaging in the inner plexiform layer at a depth of 40-50 μm visualised the dendritic arborisation of RGCs (Figure 3D). However, it can be easily seen that the SNR drops when increasing the imaging depth (for example, see Figure 6A,B).

**Figure 3.**
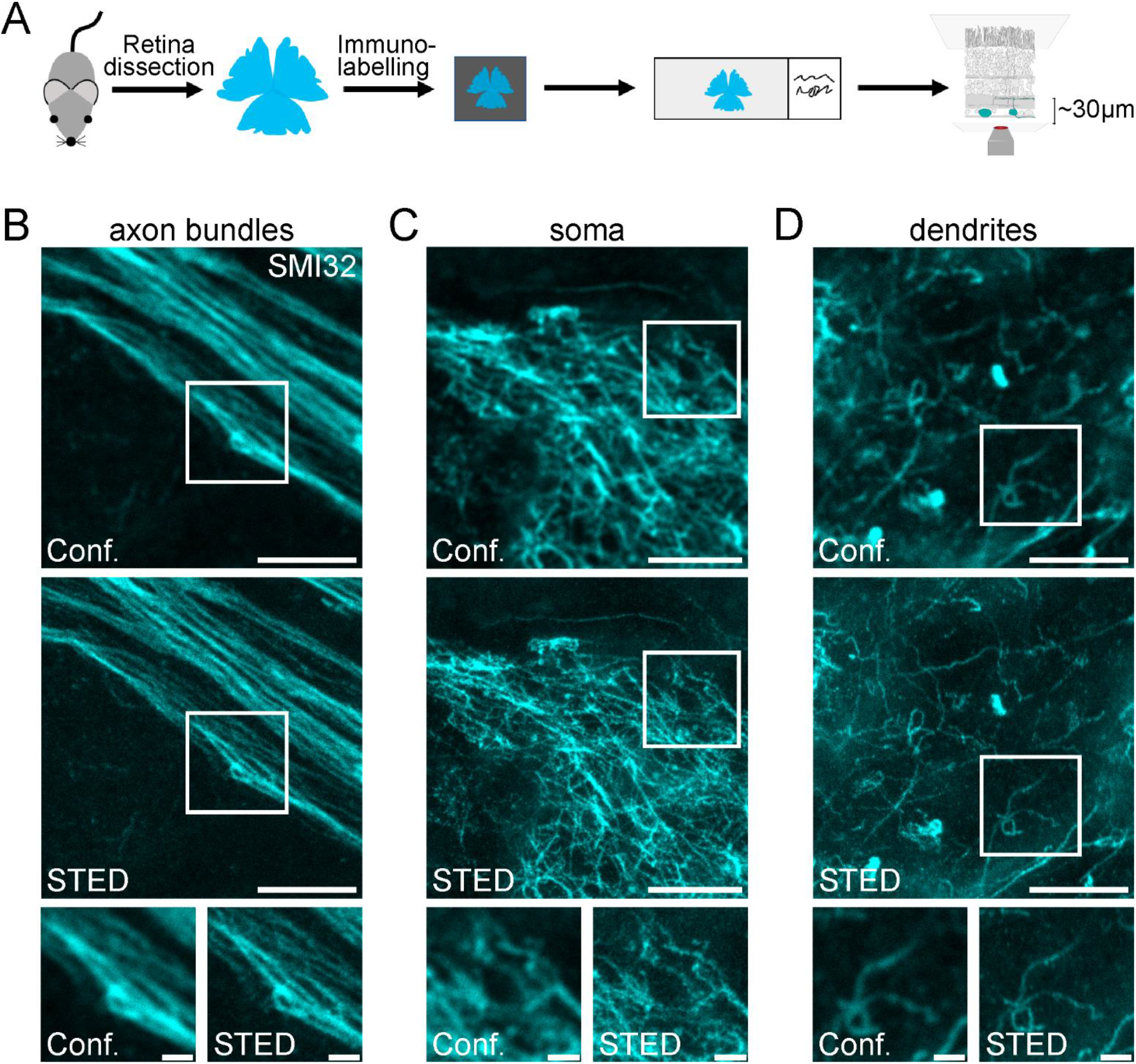
Confocal and STED imaging of SMI32-positive retinal ganglion cell structures in the inner retina. **(A)** Scheme of experimental design for imaging RGC structures in the inner retina. Retinae were dissected, mounted on filter paper, immunolabelled, mounted on glass slides and imaged. (**B-D).** Representative example images of RGCs’ axon bundles (B), soma (C) and dendrites (D) imaged in the confocal (conf.) and STED mode at a depth of ~30 μm (axons, soma) and ~50 μm (dendrites). Scale bars: 5 μm; for insets: 1 μm.

As a measure of spatial resolution, the FWHM was calculated by selecting thin filamentous structures and fitting a Gaussian curve to the intensity values of respective confocal and STED images using a custom-written R script (see Methods). The described approach allowed paired comparison between the resolution of confocal and STED images (Figure 4A). The mean FWHM of confocal images was 254.1 ± 4.7 nm, coming close to the diffraction limit of approx. 200 nm. In comparison, the FWHM of STED images was 118.2 ± 4.7 nm and therefore surpassed the theoretical diffraction limit (n = 24 structures measured, n = 3 images, mean ± SEM) (Figure 4B). As the FWHM is calculated from differently sized biological structures, it strongly depends on the structures selected and can vary between conditions. To correct for this effect, a resolution enhancement factor (REF) was calculated [29], which was defined as the ratio of STED FWHM to confocal FWHM of the same structure. Here, the REF peaked at 2.21 ± 0.07 (mean ± SEM) and ranged from 1.50 to 3.09 in the dendrites of RGCs (Figure 4C).

**Figure 4.**
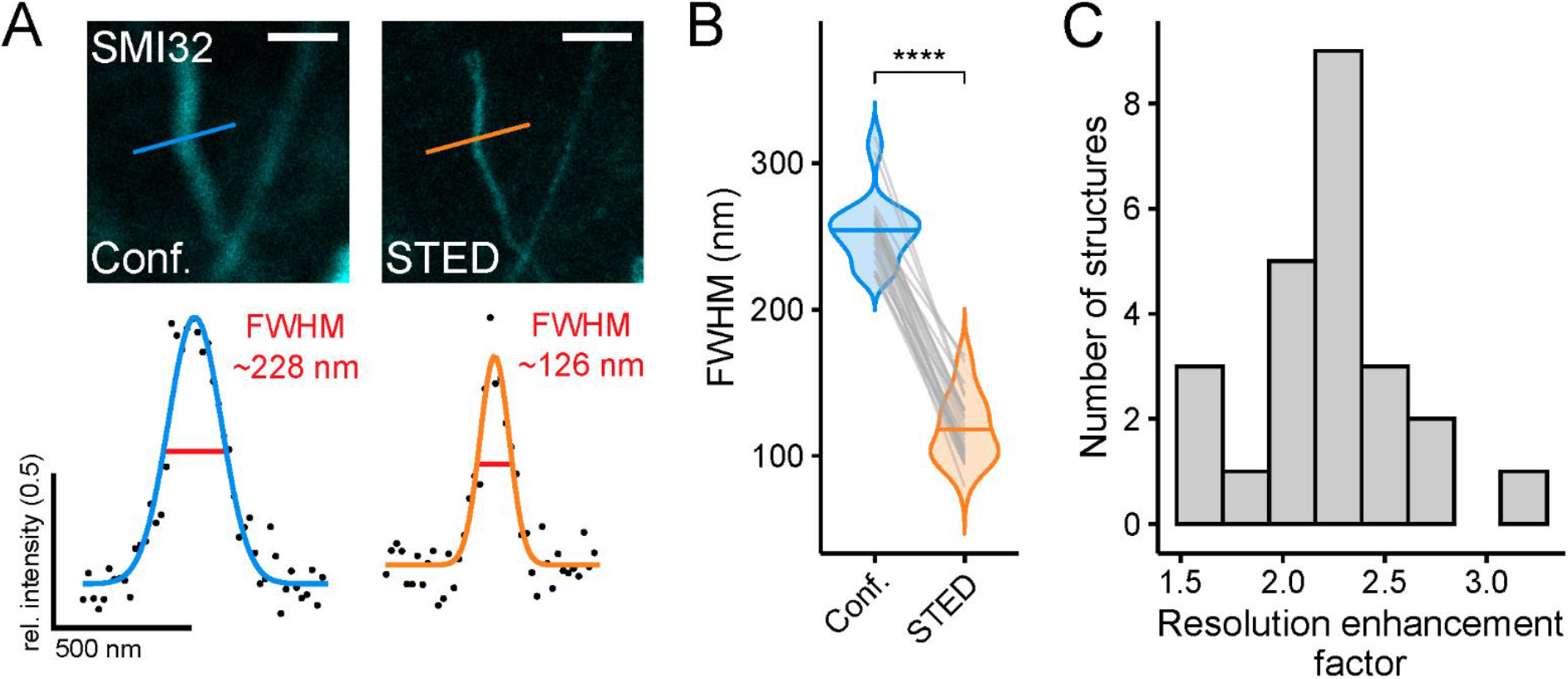
Super-resolution STED imaging of SMI32-positive retinal ganglion cell dendrites in the inner retina reveals spatial resolution enhancement. **(A)** Example structures (top, dendrite of a SMI32-positive RGC) for resolution estimation and with FWHMs for dendritic structure for confocal (blue) and STED (orange) (bottom). FWHM intensity profiles taken at the indicated position in example confocal and STED images. (**B)** Comparison of FWHMs in confocal (blue) and STED (orange) images. (p < 0.0001 = ****, Wilcoxon signed-rank test, n = 1 animal; n = 24 filamentous structures from 3 images. **(C)** Histogram of resolution enhancement factor quantified as ratio between the FHMHs of confocal and STED images (n = 24). Scale bar: 1 μm in A.

In some retinal samples, lateral and axial drift occurred at the stage of sample imaging. We tackled this problem using the Huygens Software (SVI) where appropriate. Although the image stabilisation tool performed well in lateral (xy) direction, allowing us to obtain good 3D stacks and time series, it provided no satisfying solution for axial drift (along the z-axis). The same software was used for image deconvolution (Figure 5A). We used a Classical Maximum Likelihood Estimation algorithm and adapted the program settings to obtain reliable deconvolution results. We compared both the FWHM and the signal-to-noise ratio (SNR) of STED and deconvolved STED images. While the deconvolution only marginally (though significantly) increased the resolution of STED images (STED, FWHM = 125.7 ± 7.1 nm; STED deconvolved, 107.2 ± 3.4 nm, n = 26 structures, p < 0.001, Wilcoxon signed-rank test) (Figure 5B,C), it did increase the SNR (STED, 3.3 ± 0.1; STED deconvolved, 16.2 ± 1.4, n = 26 structures, p < 0.0001, paired t-test) and smoothened intensity profiles of given structures (Figure 5B,D).

**Figure 5.**
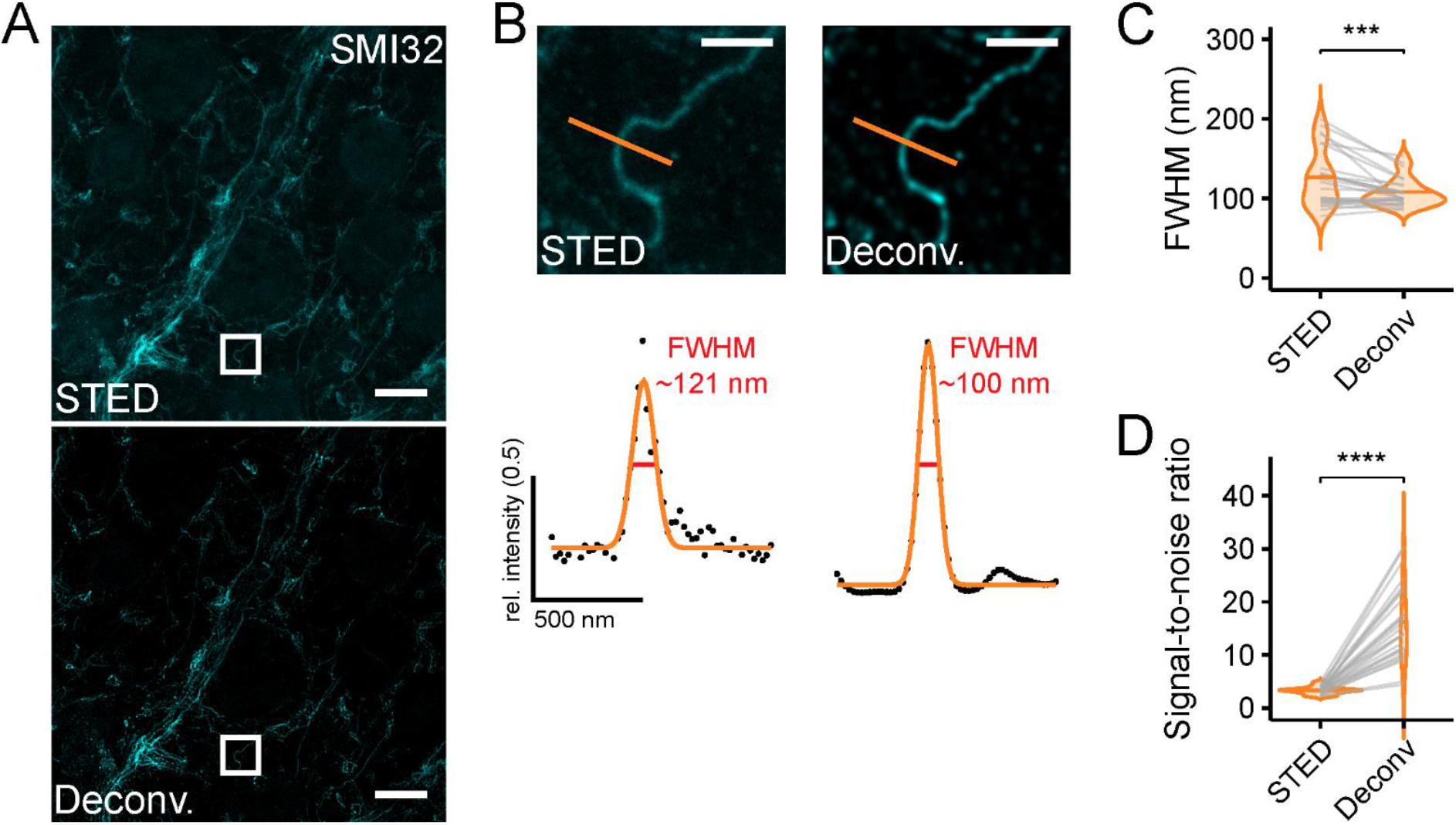
Deconvolution increases resolution and signal-to-noise ratio for retinal ganglion cell dendrites. **(A)** Representative example STED image without (top) and with deconvolution (bottom). (**B)** Top: Zoomed-in region as indicated in A. Bottom: FWHM intensity profiles taken at indicated position without (STED) and with deconvolution (Deconv.) **(C)** Quantification of the effect of deconvolution on FWHMs (p < 0.001 = ***, Wilcoxon signed-rank test, n = 1 animal; n = 26 structures from 1 image. **(D)** Quantification of the effect of deconvolution on signal-to-noise ratio (p < 0.0001 = ****, paired t-test, n = 1 animal; n = 26 structures from 1 image). Scale bars: 5 μm in A, 1 μm in B.

### Super-resolution microscopy of horizontal cell axon terminals in the outer retina

Compared with the inner retinal structures, we did not achieve an increase in resolution when we imaged SMI32-labelled HC axon terminals in the STED mode in the deep outer retina (Figure 6A,B,F,G). One possible reason for this is scattering of both the excitation and depletion laser during their passage through the tissue, and thus, misalignment of the spatially optimal laser configuration and a decrease of the STED effect. Theoretically, adaptive optics may allow the application of STED in deeper tissue while retaining super-resolution [30,31]. However, this approach would likely increase imaging duration and photo-bleaching. A more straightforward way, avoiding adaptive optics, is physical horizontal sectioning of the retinal whole-mount so that the structures of interest lay just below/at the section surface (Figure 6C, see Methods). We chose horizontal cryotome sectioning due to its widespread availability. Here, we cut off the inner layers of the whole-mounted retina and imaged (the now superficial) HC structures (Figure 6C).

**Figure 6.**
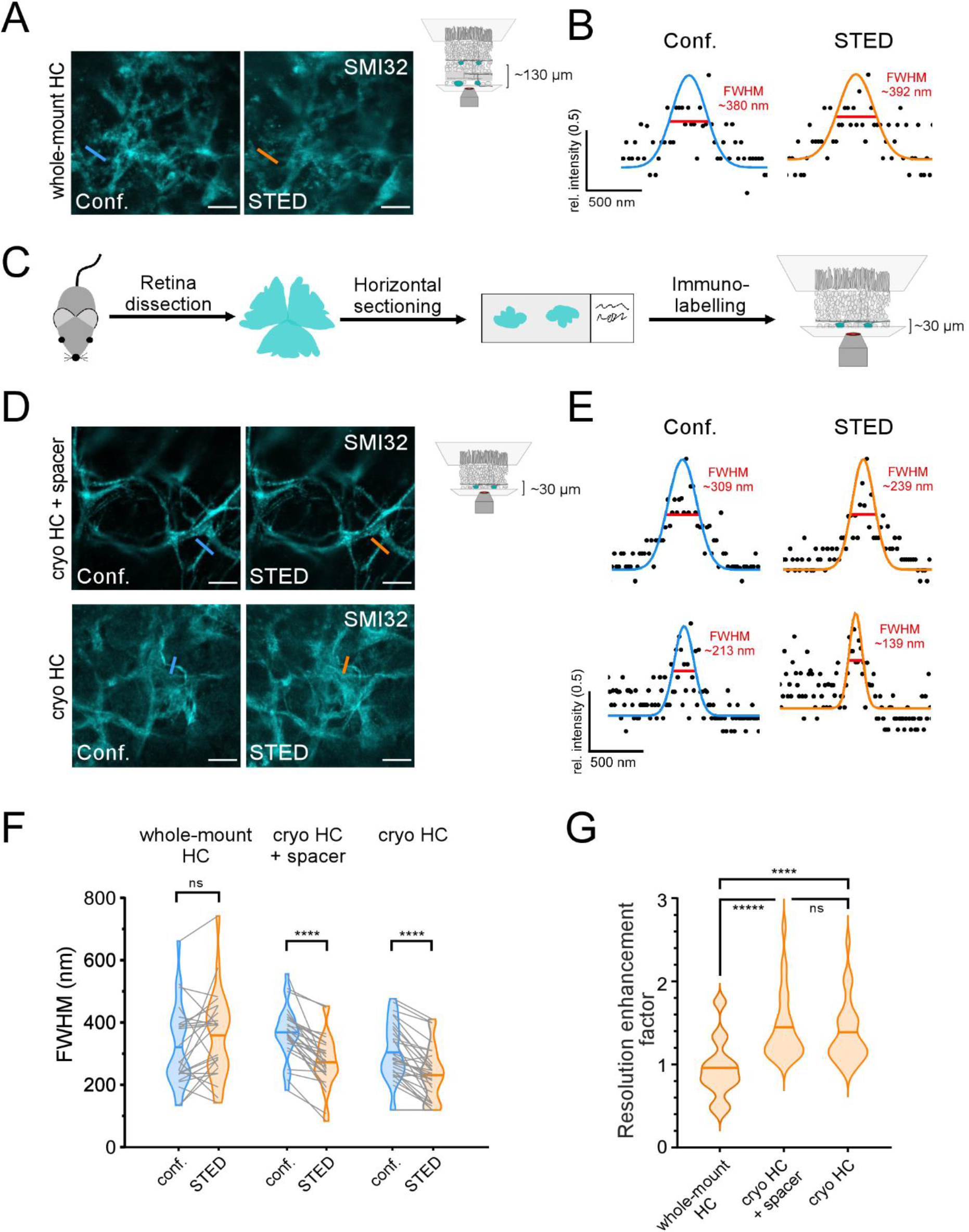
STED imaging of horizontal cell structures in the outer retina. **(A)** Comparison of confocal (left) and STED (right) images of SMI32-stained HC axon terminals at a depth of ~130 μm in the retinal whole-mount imaged through the inner retina. Each image was first taken in confocal mode and then in the STED mode to allow a direct comparison. (**B)** Confocal (blue) and STED (orange) FWHMs of the same neuritic structure as indicated in A. **(C)** Scheme of alternative experimental design for imaging structures in the deeper outer retina. Retinae were horizontally cryosectioned, mounted on glass slides, immunolabelled, and imaged. (**D)** Example confocal and STED images of SMI32-stained HC axonal structures taken with (top) and without (bottom) silicon spacers between slide and coverslip. (**E)** Confocal and STED FWHMs of the same neuritic structures in D. (**F)** Quantification of FWHMs of corresponding HC structures in confocal (blue) and STED (orange) imaging mode for whole-mount condition (whole-mount HC) and horizontally cryosectioned retina (cryo HC) with and without spacers (whole-mount HC n = 1 animal / 24 structures, Cryo HC + spacer n = 1 animal / 26 structures, Cryo HC n = 1 animal / 30 structures (p < 0.0001 = ****, ns = not significant, paired t-tests, horizontal bars indicate means). (**G)** Violin plot showing the resolution enhancement factor calculated as the ratio of the corresponding FWHMs in confocal and STED images for the three experimental conditions, horizontal bars indicate means (p < 0.0001 = ****, p < 0.00001 = *****, ns = not significant, Wilcoxon rank sum test). Scale bars: 5 μm in A,D.

Intact and horizontally cryosectioned whole-mounted retinae were stained against SMI32 using ATTO532 as the fluorophore. One set of horizontal cryosections was surrounded by a 120 μm thick silicon spacer to minimise the physical pressure of the cover slip on the retina and to avoid squeezing, while in another set, the retinal sections were directly touching the cover slip without any spacer (Figure 6D,E). Images were taken in each condition first using confocal mode and then switching to STED, and thus, imaging the exact same region. Again, we calculated the FWHMs of filamentous structures as a measure for the resolution in both confocal and STED images (Figure 6B,E). We did not find any significant change of the FWHM for STED compared to confocal images of HC structures in the intact retina (confocal, 319.5 ± 26.9 nm; STED, 358.3 ± 28.4 nm; p = 0.11, paired t-test) (Figure 6A,B,F). Interestingly, in the intact whole-mount, the variability of FWHMs for STED images was strongly increased at the HC level (Figure 6F), likely illustrating the random effect of scattering in inhomogeneous and deep tissue. In contrast, in both cryosection conditions (with and without spacers), we did not see such a pronounced effect on the variability. However, here we found significantly improved STED FWHMs with spacer (confocal 367.4 ± 16.9 nm; STED, 271.6 ± 16.8 nm; p < 0.0001, paired t-test) and without spacer (confocal, 302.7 ± 17.2 nm; STED, 230.1 ± 15.6 nm; p < 0.0001, paired t-test), thus indicating that horizontal sectioning indeed reduces aberrations in STED microscopy (Figure 6D-F). While not significantly different (confocal p = 0.09 and STED p = 0.44, 2-way ANOVA test), both confocal and STED FWHMs tended to be slightly smaller in cryosections mounted without silicon spacers, possibly reflecting a better fit with the working distance of the microscope objective. To correct for differently sized biological structures, we calculated the REF in all conditions (Figure 6G), which validated the previous results: STED in whole-mounted retinae did not alter the resolution in HC axon terminals (0.95 ± 0.07). In contrast, the average relative resolutions in the cryosectioned groups were improved by the factors 1.44 ± 0.08 (with spacer) and 1.38 ± 0.07 (without spacer). The REF in HC structures in cryosectioned retinae did not significantly differ from the whole-mounted RGC axons imaged in the same set of experiments (REF: 1.41 ± 0.10, n = 1 animal, n = 23 RGC structures; p = 0.46 compared to HC with spacer and p = 0.77 compared to HC without spacer, Wilcoxon rank sum test).

### Identification of dendritic bulbs of horizontal cells

After confirming that the antibody staining and STED imaging protocols work as anticipated for deep retinal imaging, we applied our approach to the outer circuits of the mouse retina. Bulb structures on HC dendrites have been recently identified as putative synaptic sites between HCs and BCs [27]. Therefore, we aimed at investigating this synapse type using super-resolution microscopy. We first identified HC bulb structures in the cryosectioned retina using confocal microscopy. The calcium-binding protein Calbindin is expressed throughout HCs, with anti-Calbindin staining being used to visualise the whole cell including soma, dendrites, and axon terminals. In HCs, neurofilaments are only expressed in the axon terminal system and can thus be used to distinguish between dendritic and axonal structures [32]. GABA ρ2 is a subunit of GABA_C_ receptors, which in the outer retina is exclusively expressed on BCs, thus indicating postsynaptic BC sites. Before performing a triple staining, primary antibody functionality was tested in single and double stainings, observing structures known to express the targets of the antibodies with confocal microscopy (Calbindin for HC somata, dendrites and axons, SMI32 for HC axons, GABA ρ2 for GABA ρ2-expressing receptor clusters in the HC layer).

Bulbs were characterised as Calbindin-positive and SMI32-negative structures that possibly co-localize with GABA ρ2. Bulb identification was performed using lower magnification confocal microscopy and imaging Calbindin and SMI32 in large z-stacks, spanning the whole HC layer. Calbindin and SMI32 images were superimposed, and single z-slices were manually scanned for bulb structures (see examples in Figure 7A). Surprisingly, only few SMI32-negative structures could be observed, suggesting that HC dendrites are not isolated from axons but co-fasciculate, and thus, cannot be easily distinguished under the microscope. However, several round and ‘blobby’ SMI32-negative thickenings could still be detected emerging from double positive structures, which we assumed to be HC dendritic bulbs (Figure 7A). Such identified bulbs were further examined by extracting normalised SMI32 and Calbindin intensity values along a line selection which was put through each bulb (Figure 7B,C). Bulbs were more positive for Calbindin than for SMI32 and underlying SMI32 signals did not follow the shape of calbindin signals (Figure 7B). Intensity values were also extracted from line selections of control structures (dendritic thickenings, non-bulbs), which we assumed to be double-positive (Figure 7A top panel, C). For further comparison and statistical analysis, mean staining intensities along the lines were calculated for each structure. SMI32 and Calbindin signals did strongly correlate in many non-bulb controls but never in the bulbs (Figure 7D). Indeed, the average intensity difference was significantly higher for the HC bulbs than for the non-bulb structures (bulbs, 102.51 ± 5.83; non-bulbs, −12.51 ± 10.19; p < 0.000001, Wilcoxon rank sum test) (Figure 7E). Thus, although SMI32-positive/Calbindin-positive axons and SMI32-negative/Calbindin-positive dendrites of HCs did strongly co-fasciculate, the reliable identification of dendritic bulbs was possible.

**Figure 7.**
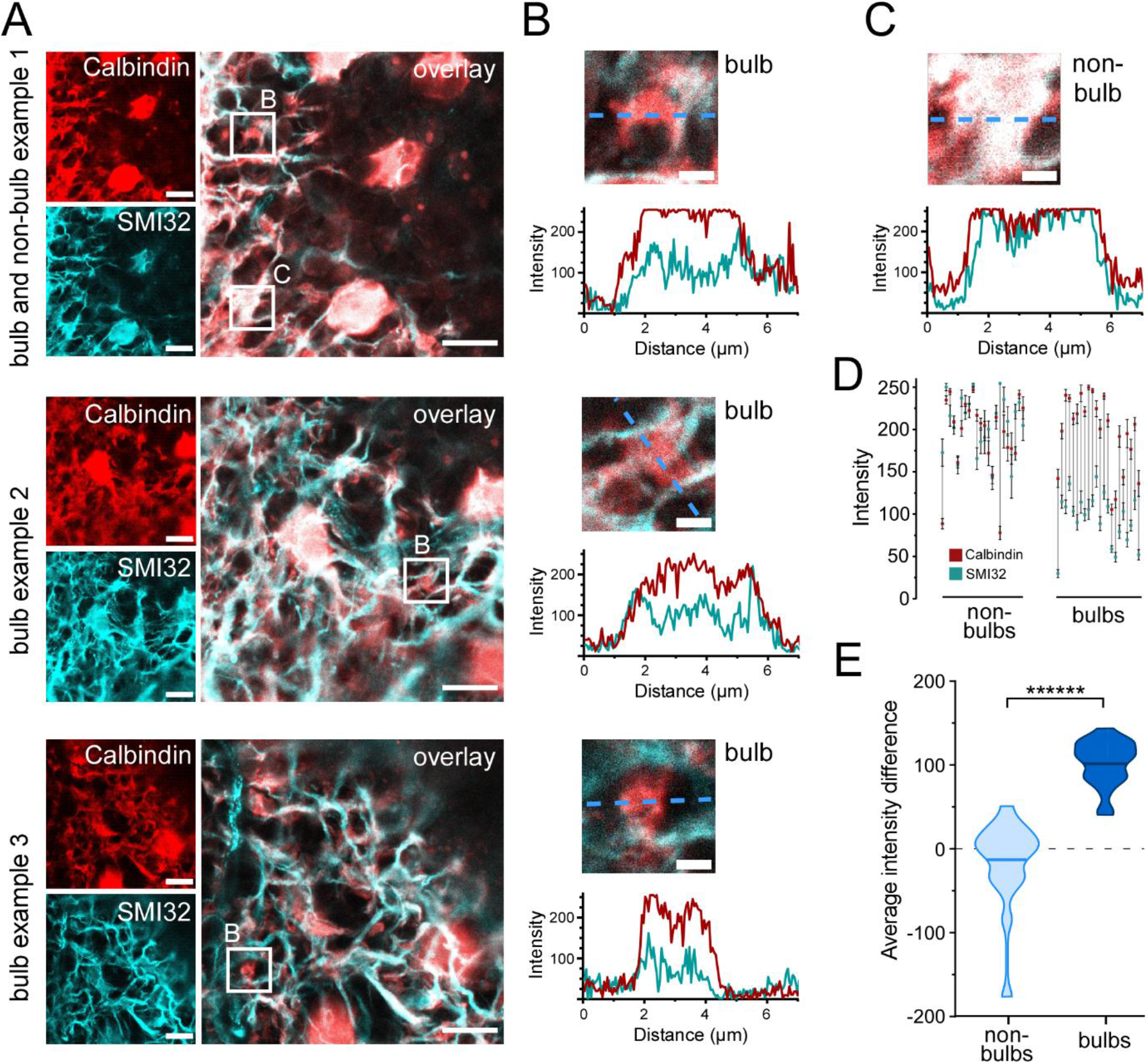
Identification of dendritic horizontal cell bulbs. **(A)** Three example images showing three HC bulbs and one non-bulb identified in confocal single plane images z-stack slices. Retinal sections were imaged for Calbindin and SMI32. Bulb structures (bulb) positive for only Calbindin and non-bulb structure (non-bulb) positive for both SMI32 and Calbindin were identified (small white squares). **(B)** Intensity values for Calbindin and SMI32 stainings were extracted along a line through the three bulbs taken from A. Plots show Calbindin and SMI32 intensity distribution across the three bulbs (dashed blue lines). **(C)** Intensity values for Calbindin and SMI32 stainings of a dendritic non-bulb taken from A. (top panel). Plot shows Calbindin and SMI32 intensity distribution across the non-bulb (blue dashed line). (**D)** Mean intensities of Calbindin (red) and SMI32 (cyan) stainings along the lines through individual non-bulbs (left) and bulbs (right). Error bars indicate 95% confidence interval. (**E)** Violin plot showing the average intensity difference. Difference per pixel was calculated for multiple bulbs and non-bulb control structures, horizontal bars indicate means (n = 1 animal, n = 22 structures for bulbs, n = 22 structures for non-bulbs, p < 0.000001 = ******, Wilcoxon rank sum test). Scale bars: 10 μm in A, 2 μm in B,C.

**Figure 8.**
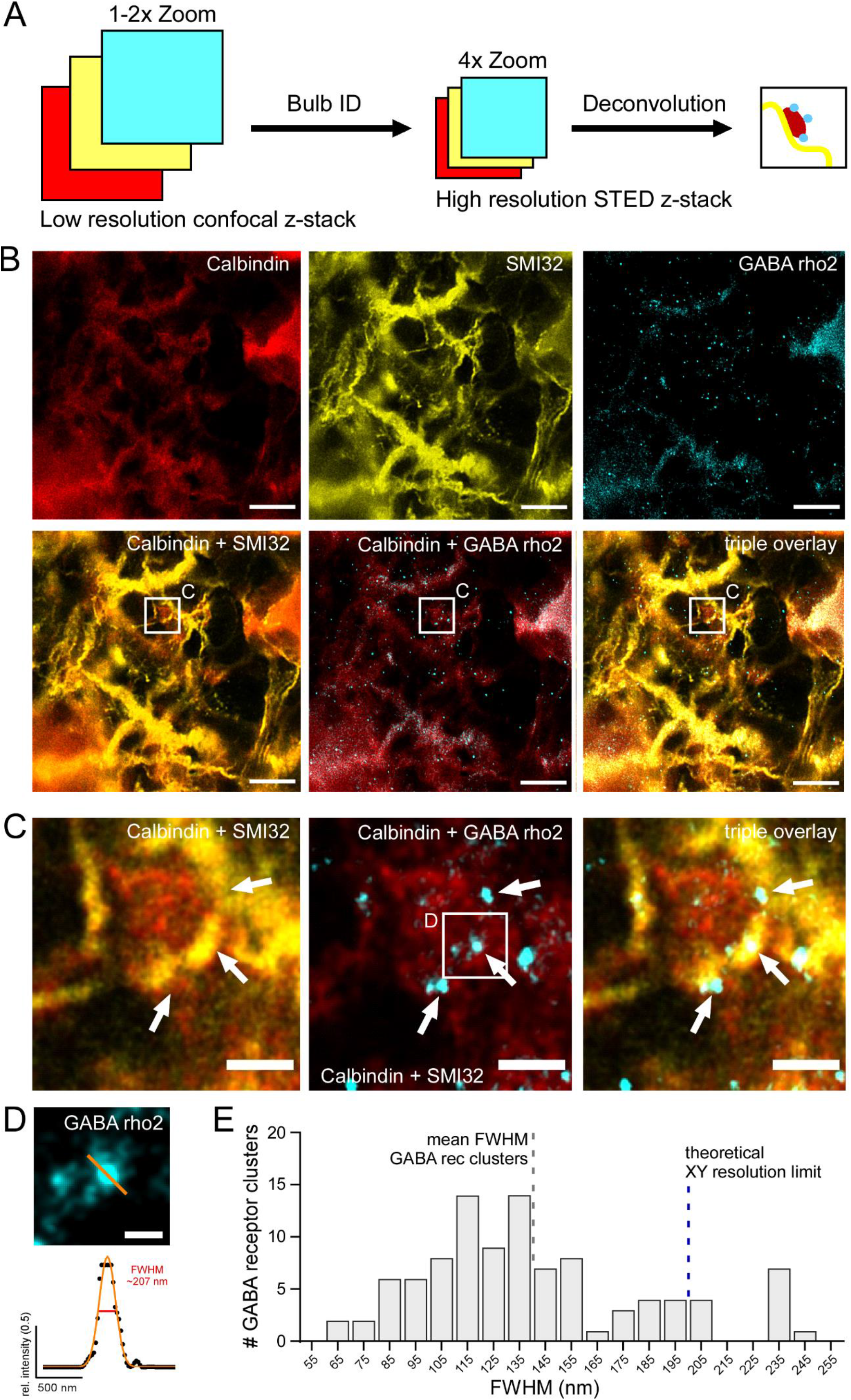
GABA_C_ receptor clusters can be localised on horizontal cell dendritic bulbs with high-resolution STED. **(A)** Schematic showing experimental design to image dendritic HC bulbs with high-resolution STED microscopy. Individual bulbs are identified in sections of large confocal z-stacks (Bulb ID, see also Figure 7). Bulbs were zoomed into, imaged as STED z-stacks and visualised using maximum z-projections and deconvolution. (**B)** Triple staining experiment against Calbindin (red), SMI32 (yellow) and GABA ρ 2 (cyan) showing a dendritic bulb (white square) imaged in the STED mode. The centre of the bulb is negative for SMI32 but positive for calbindin labelling. White square indicates the position of the bulb depicted as close-ups in C. **(C)** Zoomed-in bulb with triple staining against Calbindin, SMI32 and GABA ρ 2 showing a HC dendritic bulb (from white square in B) imaged in STED mode and deconvolved. Note that GABA ρ 2 receptor clusters (arrows) are located at the edges of bulb. (**D)** Example ρ 2-positive GABA receptor cluster (top, small white square from C) with Gaussian fit and FWHM (bottom). FWHM intensity profile is taken at the indicated position (orange line) Scale bar: 0.25 μm. (**E)** Histogram showing distribution of FWHMs of GABA ρ 2 clusters (in 10 nm bins) (n = 1 animal, n = 100 clusters). Dashed blue bar indicates xy resolution limit for confocal light microscopy, dashed grey bar shows FWHM mean for GABA receptor clusters. Scale bars: 5 μm in B, 1 μm in C.

### Identification of putative GABAergic synapse at bulb sites

To image HC bulbs and GABA ρ2 receptors with higher resolution, STED images of identified bulb structures were acquired. Bulbs were identified as described above using large confocal z-stacks, which were screened for bulbs directly at the microscope (Figure 8A). For this triple labelling approach, each primary antibody was paired with multiple secondary antibodies, coupled to different fluorophores, and imaged using both confocal and STED microscopy to identify fluorophores working optimally with three-colour STED (Figure 8B). Fluorophores of the Alexa Fluor family were tested but found to easily bleach. In the end, fluorophores of the STAR and ATTO families were chosen for their superior photostability, although in confocal imaging, they can be dimmer compared to Alexa Fluor dyes. To avoid spectral overlap in the triple staining and make use of the available excitation and depletion lasers, STAR488 was chosen for the Calbindin, ATTO532 for SMI32, and ATTO633 for GABA ρ2 staining. Bulbs were then zoomed-in using confocal live-view before Calbindin, SMI32 and GABA ρ2 stainings were subsequently imaged with the STED mode. Deconvolution was applied afterwards to increase SNR and better resolve the fine GABA receptor clusters. Interestingly, bulbs identified in confocal mode were often only weakly visible with higher-magnification STED microscopy. However, bulbs that could be observed with STED correlated with multiple GABA receptor blobs/clusters (Figure 8C). Finally, we determined the FWHMs of GABA receptor clusters in deconvolved STED images (Figure 8D). Our quantification showed that the vast majority (93 out of 100) of analysed GABA ρ2-positive clusters had an FWHM under 200 nm, surpassing the xy resolution limit of confocal light microscopy. In fact, many FWHMs peaked at 110 to 120 nm (mean ± SEM: 140.1 ± 4.4 nm, n = 100 receptor clusters), which is in the resolution limit of super-resolution STED microscopy (Figure 8E).

In conclusion, we were able to reliably identify GABA receptor clusters on HC dendritic bulbs using super-resolution STED microscopy and could show that the FWHM of the GABA receptor clusters is clearly under the resolution limit of confocal microscopy.

## Discussion

Super-resolution imaging of thick specimens has been established for different brain structures [13,33]. Here, we developed a reliable protocol for STED imaging in the whole-mount mouse retina preparation. To our knowledge, no comparable protocols existed so far. To optimise the imaging procedure, we adapted sample preparation and microscope settings. As the first retinal model system, we labelled neurofilaments in ganglion cells and found that – at a depth of up to ~30 μm – the spatial xy resolution could be increased by a factor of > 2. Next, we aimed at imaging synaptic structures of HCs in the outer retina at a depth of ~130 μm. Due to scattering of the laser light and misalignment of the excitation and depletion laser beams, super-resolution imaging in deeper layers of the outer retina did not yield resolution improvements. However, we established a method to circumvent this problem and to increase the resolution of STED microscopy when imaging in those deeper retinal layers: horizontal cryosectioning and removal of the inner retinal tissue allowed us to access HC neurites with STED imaging. Here, we visualised dendritic bulbs, which likely represent a novel type of synapse capable of GABAergic feedforward signalling from HCs to BCs. Thus, imaging in the whole-mount retina can help to describe the protein composition and scaffold at retinal synapses. Taken together, we are convinced that our protocol further expands the application portfolio of STED microscopy.

### Resolution increment is determined by the saturation factor of the depletion laser

For single-colour STED microscopy and for imaging the fine structures as GABA receptor clusters, we chose the 775 nm depletion beam because with it we obtained a higher saturation factor than with the 592 nm laser. The two lasers differed in both laser architecture and temporal properties of excitation, depletion and detection. The 775 nm pulsed/gated laser allowed us to achieve a saturation factor of 30, which was theoretically sufficient for resolution scaling up to 3 times if compared with confocal microscopy [29]. The main constraint with increasing depletion laser intensity was specimen overheating. Another problem that we faced was the loss of SNR with increasing imaging depth and light scattering in the tissue. To our knowledge, it is a common problem with several possible solutions [15,16]. We tried deconvolution and obtained a pronounced SNR improvement. To further increase contrast and improve the axial resolution, one can use a 3D STED approach with additional donut-shaped illumination perpendicularly to the optical axis [34] or adaptive optics [30,31], or alternatively horizontal cryosectioning (see below).

### Optimal resolution with the conventional antibody approach in the whole-mounted retina

A very crucial factor of every super-resolution microscopy approach is resolution estimation. For this purpose, we fitted a Gaussian to the line profiles of filamentous structures using the NLS algorithm. From the Gaussian fits, we determined the FWHMs and compared the values between imaging conditions. This approach, although widely used [12,35,36], is prone to errors [15] and relatively laborious. It requires isolated filamentous or punctuated nanometre-sized structures, which are not necessarily easy to find even in the samples labelled against cytoskeletal or calcium-binding proteins. Given the abundance of the antigen and the antibodies concentration, we had sufficient labelling density. However, given that the intermediate filament has a width of ~10 nm and is labelled by indirect immunodetection with full-sized antibodies having size of ~20 nm, the best accessible resolution is restricted to approximately 50 nm in our case [37]. Furthermore, the labelling with full-sized antibodies may introduce artefacts into the imaging [34] making some densely packed antigens inaccessible for labelling. This challenge can be overcome by targeted labelling with small molecules (e.g. nanobodies, protein/peptide-directed labelling, aptamers or click chemistry-based labelling of single amino acids [38–41].

### Sample preparation and immunolabeling for STED microscopy in the whole-mount mouse retina

In combination with the cryosection, a protocol for an immunocytochemistry triple staining for STED imaging was developed and tested in both whole-mount retinae and horizontal retinal cryosections. Antibodies against Calbindin, SMI32, and GABA ρ2 were chosen for assessing HC synapses, however, we expect that the protocol also works with other antibodies. In general, all tested antibody stainings functioned as expected, although problems with the GABA ρ2 antibody such as insufficient penetration of the tissue occurred from time to time. Possible explanations might be a lower affinity of the antibody, or more generally, in the antigen properties of GABA_C_ receptors, whose epitopes may be hidden within the double lipid membrane or beneath other synaptic proteins. Additionally, GABA receptor clusters are smaller and sparser than SMI32 and Calbindin complexes, resulting in a lower overall number of target epitopes. As Calbindin is present throughout the HC cytosol, its staining often appears to be diffuse and HC structures appear blurry, especially with higher magnification. Alternative approaches to stain whole HCs, such as immunostaining against GFP in transgenic animals or direct injection of fluorophores into single HCs, may result in signals with a better SNR [42].

### Retinal cryosectioning is compatible with standard immunocytochemistry and STED super-resolution microscopy

One major challenge of our approach was to show whether cryosectioning of the retina is still compatible with standard triple staining protocols using STED-compatible fluorophores. Here, we demonstrate the feasibility of performing complex super-resolution STED experiments in retinal cryosections. The newly developed protocol was applied to study HC dendritic bulbs, which likely represent a recently identified synapse for GABAergic feedforward signalling from HCs to BCs [27].

As STED microscopy is based on confocal microscopy technology, sharing the pinhole, it is intrinsically capable of optical sectioning, and thus, should be theoretically able to image in deeper planes of thick specimens. However, the spatial resolution of STED microscopy strongly depends on the exact alignment of the depletion laser donut around the excitation laser beam, which can be disturbed by scattering in biological samples. Multiple adaptive optics approaches, which measure and compensate for the distortions for each point, have been developed but involve multiple drawbacks, mainly high costs and increased imaging time and photo-bleaching. The method described here averts optical distortions by removing biological tissue above the plane of interest. Additionally, it enables the application of conventional STED microscopy without requiring special settings or computations and thus decreases costs, effort, and imaging time compared to adaptive optics. Cryosectioning is a mature and widely used technique. Still, some problems, including freezing artefacts as well as wrinkled and displaced sections, are commonly reported and could influence tissue integrity and STED resolution [43]. Even small artefacts, hardly visible under confocal microscopes, could influence images taken with STED. Thus, precise and meticulous work is important throughout the whole freezing and cryosectioning process.

### Comparing neurofilament structures of retinal ganglion cells and horizontal cells

In this study, we used the FWHM in biological samples as an indicator for the effective resolution. We demonstrated that both the absolute STED resolution as well as the relative resolution improvement of HC structures were significantly enhanced in cryosections compared with the intact retinal whole-mount, and thus, similar to the resolution of superficial RGC axons and dendrites in the whole-mounted retinae. It should be emphasised that in our experiments, the FWHM measured for individual RGC dendrites was lower than the FWHM in co-fasciculating HC neurites. Thus, the measured FWHM does not necessarily reflect the absolute theoretical resolution of the microscope but, for instance, depends on the width of the measured structure, on the thickness of the biological specimen and the SNR. Still, the FWHM has been used as a reliable indicator for resolution in past studies [44,45]. Furthermore, the relative resolutions, which were subsequently calculated, partially correct for different-sized structures and confirm the effects observed with the absolute FWHMs. The robustness of curve fitting and thus FWHM calculation also strongly depended on the SNR of the structures selected. For example, in some STED images, the SNR was frequently inadequate, and the accuracy of the fitted curves had to be manually reviewed for each structure. Due to the low number of emitted and detected photons, a low SNR is a general issue with STED imaging. Thus, for each experiment appropriate settings must be determined and the right balance between signal and resolution has to be defined. Theoretically, STED microscopy can reach xy resolutions in the low nm range and, using fluorescent beads, PSFs with as little as 5.8 nm width have been recorded [46]. However, these resolutions cannot be currently achieved in biological samples and, while proof of concept studies could produce resolutions of around 20 nm [47], the measured maximal resolution is in practice often limited by target size, optical distortions, photo-bleaching, and labelling strength. In this study, FWHMs of below 100 nm, and thus far beyond the diffraction limit of confocal microscopy, were observed. Still, even under best possible STED imaging conditions, the FWHMs of the HC structures were on average above 200 nm. One reason for this could simply be the relatively large size of the neurofilament structures stained in HCs. This possibility is supported by the fact that although STED resolutions in this experiment are beyond the diffraction limit, they still represent a significant improvement compared to the confocal FWHMs of the same structures. Another argument in favour of this view comes from imaging individual and sparse GABA receptor clusters at HC bulbs: here, the mean FWHM is around 140 nm, and therefore, is close to the mean FWHM of RGC neurofilaments.

Huygens deconvolution was applied to STED images to further increase the resolution and improve the SNR. Deconvolution algorithms are computational methods that try to recalculate the original optical scene in the sample by subtracting effects of known optical distortions and diffractions from the image [48] and that were used with STED microscopy before with excellent results [49,50]. In our case however, only minor improvements of FWHM could be observed after deconvolution, although SNR mostly appeared to be increased. A problem with the employed deconvolution algorithm might be that it is “blind”, meaning that it uses a theoretical PSF that was calculated for the microscope setup present. A better approach would include measuring real PSFs using the exact imaging conditions, which might enable the deconvolution to predict optical aberrations more accurately.

### Horizontal cell bulbs likely represent GABAergic pre-synapses

Horizontal cells are essential for the generation of centre-surround receptive fields in BCs [51]. While most studies focused on the complex HC feedback mechanisms to photoreceptors, evidence for direct HC-to-BC synaptic contacts has been found in both non-mammalian and mammalian retinae [25,26,52–54]. HC feedforward signalling is likely GABAergic and dependent on vesicular release [55,56]. Recently, bulbs on HC dendrites have been observed in 3D electron microscopy reconstructions and identified as possible synaptic contacts, with most bulbs contacting either other HC bulbs or BC dendrites [27]. Furthermore, Behrens and colleagues demonstrated the presence of mitochondria in dendritic bulbs and used immunolabeling to reveal that bulbs of individually stained horizontal cells co-localize GABA ρ2 receptors. In this study, bulb identification was performed by using Calbindin as a marker for the entire HC and additionally labelling SMI32 to counterstain HC axon terminals. This allowed the distinction between dendrites and axons and by calculating the intensity differences between Calbindin and SMI32 signals quantitative characteristics of bulbs could be defined. We originally expected to find half of the HC structures double-positive, representing axons, and the other half only positive for Calbindin, representing dendrites. However, very few Calbindin-only-positive structures could be observed. Space limitations in the outer plexiform layer and a high density of HC structures likely result in strong co-fasciculation of HC dendrites and axons and an overlay of single- and double-positive structures. Still, by imaging large z-stacks with low-magnification in the confocal mode, single bulb structures were frequently identified. These structures typically emerged from double-positive filamentous structures and occasionally overlapped with additional SMI32 signals, which might be the result of HC axons stratifying along the bulbs. Nevertheless, these structures showed no correlation between Calbindin and SMI32 signals and were thus further considered dendritic bulbs.

By detecting the ρ2 subunit of GABA_C_ receptors, which in the retina are exclusively expressed on BCs [57], at Calbindin-positive/SMI32-negative bulbs, we could specifically investigate the role of bulbs in HC to BC GABAergic signalling. Some GABA ρ2 signals were detected on or in direct proximity to bulbs, thus indicating co-localization of bulbs with BC postsynaptic sites, and supporting the notion that bulbs are HC-to-BC synapses. Still, the presence of additional synaptic markers (e.g. presynaptic proteins) would provide further evidence for the type and mechanism of bulb synapses. For instance, synaptic proteins of the release machinery in HC dendrites as well as GABA and GAD65 have been found in mammalian HCs [27,56,58]. It might be compelling to monitor these targets with super-resolution microscopy and compare their localization within bulbs to the GABA receptors. Further intriguing imaging targets are voltage-gated calcium channels, although antibodies against them are either extremely subtype-specific and have a low efficiency or are very unspecific, resulting in cross-reactions with other targets [59]. In general, staining as many targets simultaneously as possible would enable the extraction of much more information about the synapse ultrastructure, however, immunocytochemistry gets increasingly difficult the more simultaneous stainings are applied and chances for unspecific bindings or fluorescent crosstalk grow. Functional experiments unravelling the mechanisms and function of bulb synapses would be highly desirable, but selectively monitoring or manipulating HC-to-BC synapses is challenging due to the complexity of outer retinal circuits (e.g. simultaneous feedback and forward signalling within a single interneuron) and the presence of similarly complex synapses between photoreceptors, BCs and HCs in close proximity. Nonetheless, structural super-resolution studies such as the present work might provide novel access points for further experiments.

## Acknowledgements

We thank Merle Harrer and Gordon Eske for excellent technical support, and Sylvia Bolz and Christine Henes for assistance with the cryotome. We thank Karin Dedek, Christian Puller and Bettina Kewitz for helpful discussions. We thank Dominic Gonschorek, Jonathan Oesterle, and Zhijian Zhao for critical reading of the manuscript and discussion.

